# Northward expansion of *Rugulopteryx okamurae (Dictyotales, Ochrophyta)* in the southeastern Iberian Peninsula: first record in Calpe (Comunitat Valenciana)

**DOI:** 10.64898/2026.07.07.736054

**Authors:** Elena Rodríguez García, Jaime Fernández del Campo, John Y. Dobson, Eva S. Fonfría, Cesar Bordehore, Carolina Pena-Martín

## Abstract

The non-indigenous brown macroalga *Rugulopteryx okamurae* has emerged as one of the most aggressive marine invaders in European waters, deeply altering benthic communities and causing severe socioeconomic impacts. While its expansion has been extensively documented along the southern Iberian Peninsula, understanding the dynamics of its northward range expansion along the Spanish Mediterranean coast remains critical for coastal management. This study documents the first formal record of *R. okamurae* in Calpe (Alicante), representing its current northernmost distribution limit within the Comunitat Valenciana. Sampling was conducted through an initial opportunistic scuba diving observation along the surrounding waters of the Penyal d’Ifac Natural Park, followed by targeted underwater surveys and an ad hoc inspection of commercial bottom-trawling nets drying at the port of Calpe during June 2026. Morphological and anatomical identification was confirmed through cross-sections of the thallus under optical microscopy, revealing the presence of both the thick and intermediate morphotypes of the species. The collected specimens were found either entangled within a native photophilic algal canopy in shallow waters or recovered from deeper offshore fishing grounds. Given the absence of records in the area during 2023–2025 surveys, these findings suggest either a very recent front-wave colonization event or a contribution from nearby, yet undetected, established patches, driven by secondary local dispersal mechanisms such as drifting fragments and explicitly highlighting commercial fishing activities as an active vector. Furthermore, considering that the species was recorded within a marine protected area and deeper environments, these results highlight a potential ecological threat to local benthic ecosystems, emphasizing the urgent need f or competent authorities to implement spatiotemporal monitoring and public awareness campaigns to prevent the definitive establishment of this invader.

## INTRODUCTION

The introduction of non-indigenous species (NIS) represents one of the most severe threats to biodiversity and ecosystem services of the Mediterranean Sea. Among recent macroalgal introductions, the brown seaweed *Rugulopteryx okamurae* (E.Y. Dawson) I.K. Hwang, W.J. Lee & H.S. Kim (*Dictyotales, Ochrophyta*), native to the temperate and subtropical waters of the Northwest Pacific, has emerged as one of the most aggressive marine invaders in European waters (García-Gómez *et al*., 2020).

Although *R. okamurae* was first recorded in the Mediterranean basin in 2002 within the Thau Lagoon (France), where it remained non-invasive (Verlaque *et al*., 2009), its ecological behavior shifted dramatically a decade later. In 2015, massive blooms and beachcast biomass (arribazones) were detected in the Strait of Gibraltar and Ceuta (Ocaña *et al*., 2016). Since then, the species has exhibited an exponential and devastating expansion along both the Atlantic and Mediterranean shores of the southern Iberian Peninsula and North Africa (El Madany *et al*., 2024), deeply altering infralittoral rocky reef communities from 0 to 40 meters deep due to its high vegetative propagation, chemical defenses, and ecological plasticity (García-Gómez *et al*., 2021a; Cobos-Mateo *et al*., 2024). This behavior has not only caused severe ecological disturbances but has also resulted in socioeconomic consequences, including negative impacts in tourism and fisheries-related activities (Laamraoui *et al*., 2024). As a result, *R. okamurae* is currently included in the Spanish List of Invasive Alien Species (MITECO, 2020) as well as in the List of Invasive Alien Species of Union Concern (EU, 2022).

This invasive success is largely explained by its biological traits, as *R. okamurae* exhibits a high dispersal capacity through vegetative asexual reproduction, primarily via propagules and monospores, which have facilitated the extensive colonization of large areas of rocky substrata (Altamirano *et al*., 2016). The initial establishment of the species occurs preferentially on hard, well-lit rocky substrates (García-Gómez *et al*., 2020). However, it can also colonize subtidal rocky habitats with lower light availability, particularly in sheltered environments, as well as coralligenous and pre-coralligenous communities (Sempere-Valverde *et al*., 2020; García-Gómez *et al*., 2021b).

In addition to these reproductive strategies, dispersal via detached fragments constitutes an important colonization mechanism, whereby algal masses separated from the substrate remain floating over the thalli of resident species (Rueda *et al*., 2023). Once introduced into a benthic community, *R. okamurae* can establish itself through different spatial colonization strategies. The most frequent mechanisms include lateral displacement of resident species and the occupation of space made available following the decline or death of pre-existing organisms. Another relevant strategy is epiphytic growth on native macroalgae. The species has also been observed developing on resident macroalgae without directly attaching to their thalli (García-Gómez *et al*., 2021b).

The vector of secondary dispersal of *R. okamurae* —driven by shipping, ballast waters, and recreational boating— has facilitated its expansion northward along the Spanish Mediterranean coast in recent years (García-Gómez *et al*., 2021b). After colonizing the Alborán Sea (Estévez *et al*., 2022), its presence was confirmed further east and north, reaching the Gulf of Alicante in the Comunitat Valenciana (GVA, 2023). However, understanding the exact dynamics of its northward range expansion is crucial for implementing early warning systems and containment strategies before the species fixes firmly onto local, highly vulnerable ecosystems like *Posidonia oceanica* (L.) Delile meadows or coralligenous reefs.

The ecological plasticity of *R. okamurae* is further reflected in its ability to colonize a wide variety of artificial substrates. The species has been recorded on port infrastructures, breakwaters, metallic surfaces of vessels, and diverse marine debris, including abandoned fishing nets, ropes, plastics, tires, and glass bottles (García-Gómez *et al*., 2021b). This trait significantly enhances its dispersal and establishment probabilities by exploiting anthropogenic debris as additional vectors of spread (Occhipinti-Ambrogi & Savini, 2003). Based on these opportunistic traits, predictive distribution models further suggest a high potential for expansion into new areas along the southern and eastern coasts of the Iberian Peninsula, including ecologically valuable and protected sites (Muñoz *et al*., 2019).

In line with these predictions, and following its expansion along the southern Iberian coasts, *R. okamurae* has been recorded in other areas of the Spanish Mediterranean, including northern locations in Catalonia (de Torres *et al*., 2025). In the Comunitat Valenciana, the first formal record was recently documented by Terradas-Fernández *et al*. (2023) in Alicante Bay, where it was found colonizing dead *P. oceanica* matte.

In this context of progressive regional spread, the present study documents the first record of *R. okamurae* in Calpe (Alicante), representing the current northernmost distribution limit of this species within the Comunitat Valenciana. This new record provides evidence a progressive northward expansion of the species from its previously known focus in Alicante Bay. We describe its bathymetric distribution and habitat preference in this new locality. Given that the presence was recorded within a Nature 2000 Special Area of Conservation (ESZZ16006 Espacio marino de Ifac), monitoring this geographic expansion is essential for assessing the vulnerability of local benthic communities and providing baseline data for regional coastal management.

## MATERIAL AND METHODS

### Sampling site and specimen collection

Initial samples were collected on 31 May 2026 during a recreational scuba dive along the southern face of the Penyal d’Ifac (Calpe, Alicante), at depths ranging from 5 to 8 meters (Figure 1.A-B). The samples were kept refrigerated in zip lock bags and taken to the Phycology Laboratory of the Department of Environmental Sciences and Natural Resources at the University of Alicante for subsequent examination.

**Figure 1.**
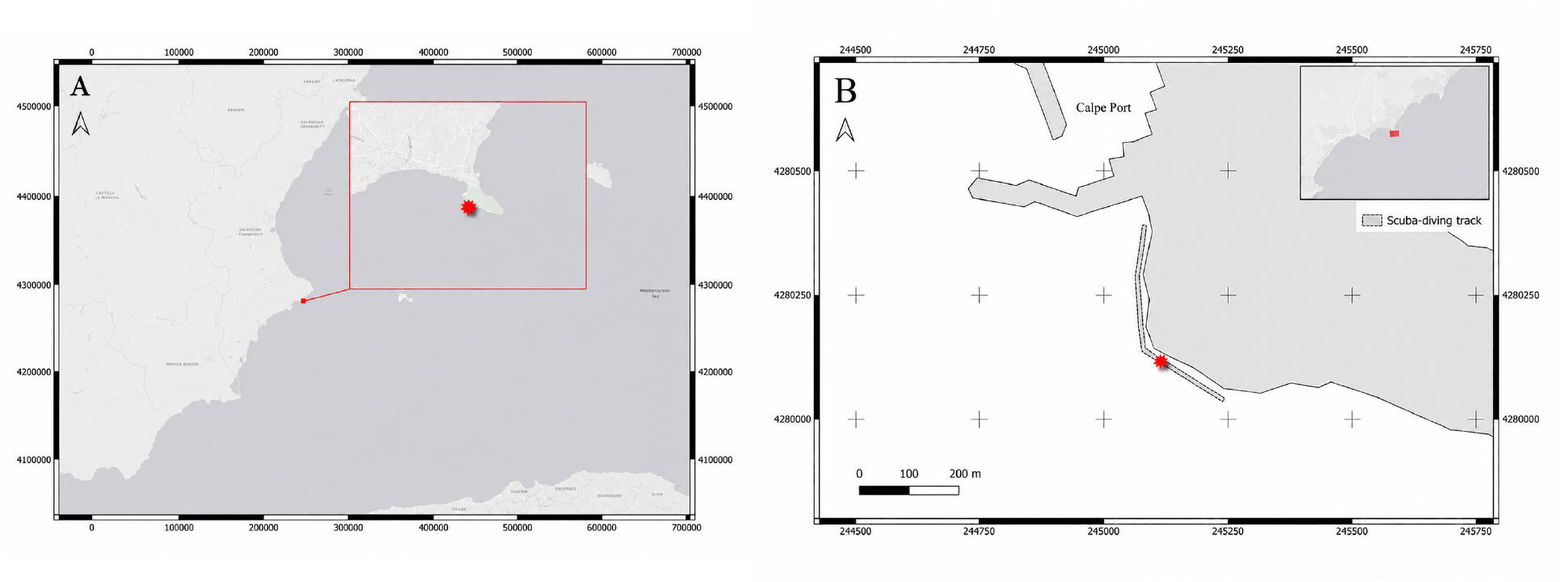
Location of the *Rugulopteryx okamurae* record. (A) Regional context within the southeastern coast of the Iberian Peninsula (Alicante, Comunitat Valenciana). (B) Detailed view of the southern face of Penyal d’Ifac, showing the specific dive track conducted during the underwater survey. Map coordinates: ETRS89 / UTM Zone 31N (EPSG:25831).

Once the identity of one of the specimens was confirmed as *Rugulopteryx okamurae*, three further dives were carried out in the area on 12, 16 and 27 June. During these surveys, the same route as in the initial dive was followed, and a systematic visual search was conducted along approximately 550 linear meters of coastline, covering an estimated 6 meter wide observation strip centered on the 5 meter depth isobath. Underwater videos were recorded using two GoPro cameras, and additional specimens with a similar appearance to *R. okamurae* were sampled and transported as described above.

Additionally, on 28 June 2026, three macroalgal specimens were recovered from the nets of commercial bottom-trawlers at the Port of Calpe. These nets were being air-dried in the sun after fishing operations. After collection, transportation to the lab was conducted as previously mentioned, to confirm their taxonomic identity.

### Morpho-anatomical characterization

The identity of the specimens was investigated in a fresh state. Transverse sections of the thalli were hand-cut with a razor blade under a binocular stereomicroscope Olympus SZX12 with objectives DFPL-1XPF and DFPL-2X. Measurements and microphotographs were then made using an Olympus SC100 digital camera mounted on an Olympus BX41 optical microscope. Based on anatomical and morphological features, taxonomic determination was performed following the specific identification keys provided by Cormaci *et al*. (2012) and in accordance with reference literature (Hwang *et al*., 2009). Subsequently, the specimen collected via scuba diving was deposited as voucher material in the Herbarium of the University of Alicante under the accession number ABH-Algae 1001, while the specimens recovered from the fishing nets were incorporated under the number ABH-Algae 1002.

## RESULTS

### Phytocenosis at the scuba diving sampling site

The benthic community at the underwater surveying site was characterized by a heterogeneous mosaic of alternating rocky reefs and sandy substrates, where scattered patches of *Posidonia oceanica* meadows were present. On the hard substrates, the assemblage corresponded to a well-developed photophilic algal community. This canopy was dominated by erect macroalgae, with a conspicuous presence of *Padina pavonica* (L.) Thivy, *Halopteris scoparia* (L.) Sauv., the alien species *Asparagopsis* sp., and several native species of the genus *Dictyota* J.V.Lamour. The understory and basal stratum were characterized by squamulose non-calcified forms such as *Peyssonnelia* sp., alongside a diverse representation of both non-geniculate and geniculate coralline algae. Notably, the single collected specimen of *Rugulopteryx okamurae* obtained via diving was found entangled among these native dictyotalean species. Consequently, it was not possible to definitively determine whether its thallus was securely attached to the rocky substrate or if it had been transported as drift material within the water column from nearby areas.

### Morpho-anatomical description

Of all the specimens collected by scuba diving, only one (sampled on 31 May 2026, at 6 m depth, 38°37’58.12”N 0°4’19.38”E) was identified as *R. okamurae*, while all the others belonged to the genus *Dictyota*. Collected specimen of *R. okamurae* reached up to 7 cm in height. Thallus width ranged from 2 mm near the basal region up to 5 mm in its widest central sections (Fig. 2.A). Thalli exhibited a yellowish-brown to olive-green coloration. Branching was irregularly and anisotropously dichotomous, displaying up to 10 divisions with obtuse apices. Neither gametangia, sporangia, nor propagules were recorded in the analyzed material. Cross-sections performed in the middle section of the thallus revealed a single-layered (monostromatic) medulla at the center, showing 3–4 cell layers at the margins (Fig. 2.B). This diagnostic internal anatomy allowed a definitive morphological confirmation of the species. Additionally, the specimen’s morphology corresponds to the thick morphotype of the species (Sun *et al*., 2006; Salido & Altamirano, 2020).

**Figure 2.**
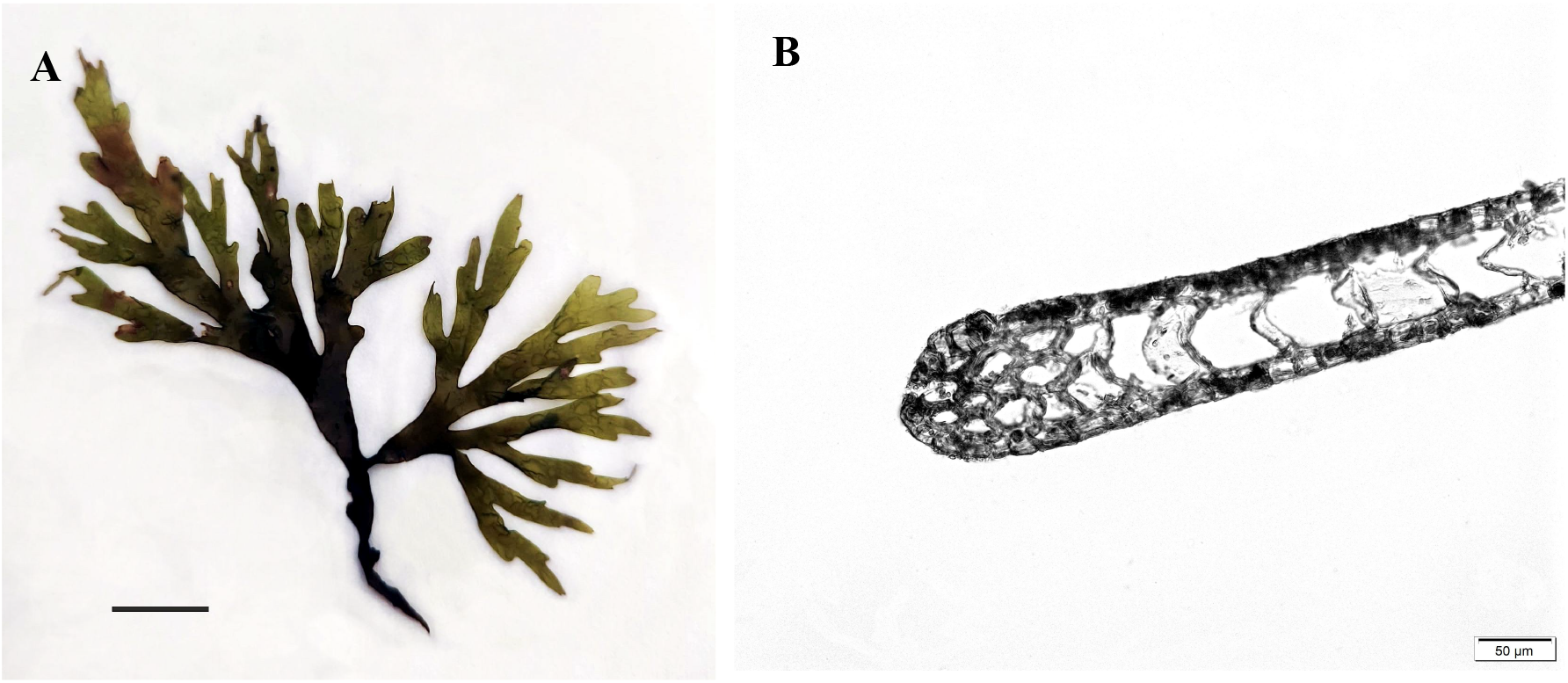
Morphological and anatomical features of the collected *Rugulopteryx okamurae* specimen from Calpe (Alicante, Spain; voucher ABH-Algae 1001). (A) General habit; scale bar = 1 cm. (B) Cross-section of the middle thallus showing the characteristic internal anatomy.

The specimens obtained from the bottom-trawlers were also taxonomically confirmed as *R. okamurae*, exhibiting heights between 2 and 5 cm, and a highly variable thallus width ranging from 1 mm near the apices up to 5 mm in the central sections of the widest individuals (Fig. 3.A). Morpho-anatomical examination revealed features characteristic of the intermediate morphotype (Sun *et al*., 2006; Salido & Altamirano, 2020): the basal and middle zones of the thallus were wide, exhibiting a medulla that was monostromatic at the center but multi-layered near the margins (Fig. 3.B), whereas the apical sections showed a distinct narrowing and a simplified internal structure.

**Figure 3.**
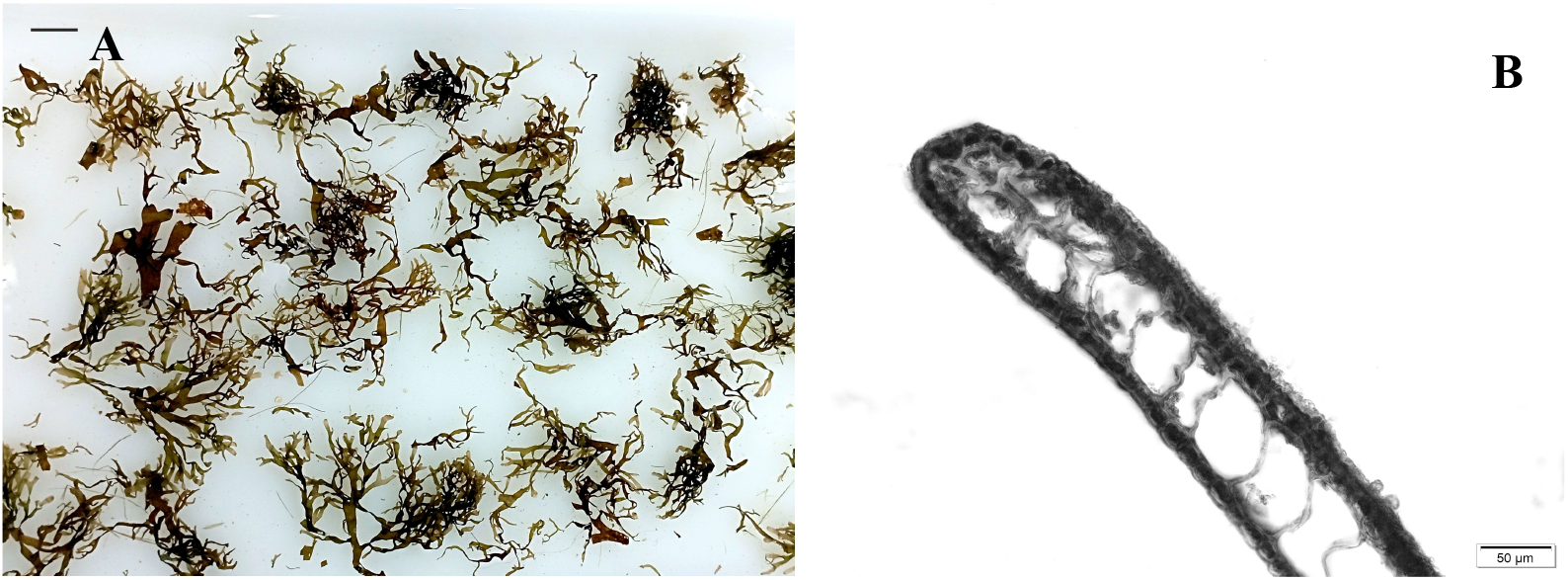
*Rugulopteryx okamurae* specimens from bottom-trawling nets. (A) General habit of the collected specimens. Scale bar = 1 cm. (B) Cross-section of the middle zone of the thallus, showing a central monostromatic medulla that becomes multi-layered at the margin.

## DISCUSSION

Although only one specimen was found *in situ* during the underwater surveys —with all other subaquatic samples consisting exclusively of native *Dictyota* species— both this single shallow individual and the specimens later recovered from the fishing port are undoubtedly *Rugulopteryx okamurae*. The macroscopic features of the initial collected specimen further support its identification within the context of the region’s invading populations. The observed gross morphology of this individual aligns consistently with the expected seasonal thick morphotype described for *R. okamurae* in south-Iberian waters during this period of the year. This species is well-known for exhibiting notable temporal morphological plasticity, adjusting parameters such as thallus thickness, width, and branching density as an adaptive response to shifting environmental and seasonal fluctuations (Salido & Altamirano, 2020). The fact that this particular individual matches these localized phenological traits reinforces that it represents a typical vegetative representative of the invading front currently expanding along the coast.

The finding of this first individual within a shallow, photophilic algal community, characterized by the presence of native Dictyotales and the alien species *Asparagopsis* sp., is highly consistent with the ecological preferences reported for this invader during its initial colonization stages in the Mediterranean. Similar shallow macrophytic assemblages and rocky habitats have been identified as highly vulnerable areas where *R. okamurae* successfully establishes (García-Gómez *et al*., 2020; Marletta *et al*., 2024). Furthermore, the fact that the specimen was found trapped within the canopy of other macroalgae supports the hypothesis that detached, drifting fragments play a crucial role in the secondary local dispersal and front-wave colonization of new coastal sectors, a mechanism also highlighted during its recent expansion in other Mediterranean areas (García-Gómez *et al*., 2020, 2021b; Marletta *et al*., 2024). Consequently, the presence of this single shallow specimen could either represent the very first front-wave settlement in Calpe or, alternatively, indicate the existence of small, yet undetected, established donor patches in nearby coastal sectors from which this fragment detached. Therefore, rather than an isolated event, the presence of this invader in Calpe likely reflects the typical front-wave signature of a progressive northward range expansion, exploiting receptive native communities.

Several studies and predictive models have forecast the expansion of this species along the Spanish coastline (Muñoz *et al*., 2019; Sainz-Villegas *et al*., 2022). However, to the best of our knowledge, this study represents the first record reported between two major port areas where the species has been detected: Alicante and Barcelona (Terradas-Fernández *et al*., 2023; de Torres *et al*., 2025). Both locations are characterized by intense maritime traffic and, as a result, they receive high volumes of ballast water, possible introduction pathway of this species (Terradas-Fernández *et al*., 2023). Although the mechanism by which *R. okamurae* reached Calpe has not yet been established, ballast water can reasonably be ruled out, given that Calpe port does not receive large cargo ships or any ship large enough to exchange ballast waters. Other port-related activities, such as fishing (e.g. the potential transport of algal fragments entangled in fishing gears), as well as natural dispersal through marine currents, cannot be excluded. In this context, the recovery of additional *R. okamurae* specimens from bottom-trawling nets at the Port of Calpe on 28 June 2026 highlights two critical aspects of this regional invasion. First, the presence of the intermediate morphotype—combining a robust basal structure (with multi-layered margins) with narrow apical sections—reflects the complex morphological plasticity of the species during the summer period in the western Mediterranean. Since the trawler from which the nets were inspected has recently been operating north of Calpe Port, specifically within Moraira Bay facing Cap d’Or (see Figure 4.B) at a depth of approximately 50 m, this finding strongly implies that the invasive alga is already colonizing deeper marine habitats, expanding its vertical distribution beyond the shallow rocky reefs surveyed by diving. Second, it confirms commercial fishing activities as a major vector for secondary dispersal. Trawling nets can inadvertently fragment and transport biomass; if these nets are cleared or deployed in uninvaded areas, they may facilitate the establishment of new focus populations along the coast.

**Figure 4.**
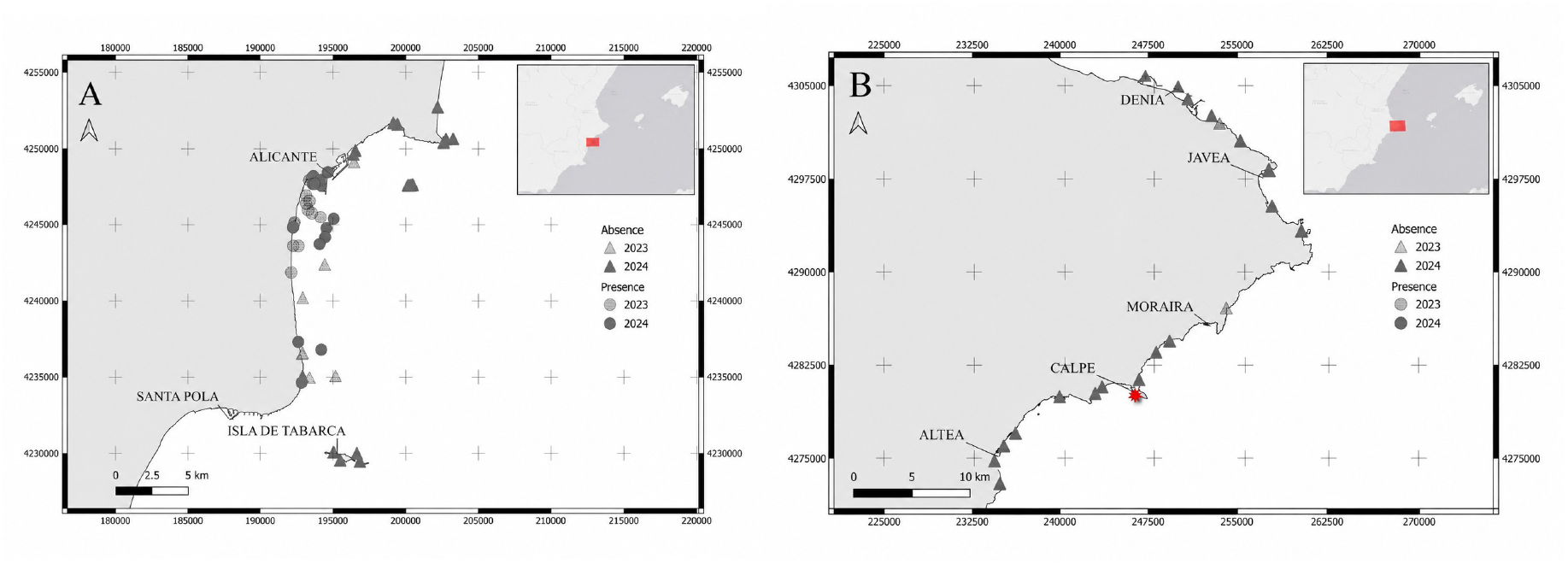
Presence/absence of *Rugulopteryx okamurae* during the 2023–2024 period. (A) Distribution across the Alicante province coastline. (B) Detailed view of the Marina Alta region. Data source: Regional Government of Valencia (GVA) and original sampling. Map coordinates: ETRS89 / UTM Zone 31N (EPSG:25831).

The date of introduction in the Penyal de Ifac area also remains uncertain. According to data from the regional environmental authority (Conselleria de Medi Ambient, Infraestructures, Territori i de la Recuperació, Generalitat Valenciana) and information from our own surveys conducted in Dénia and Jávea during 2023–2024, the presence of *R. okamurae* appears to be restricted to the area surrounding Alicante, not extending beyond Cabo de las Huertas as its northern limit (Figures 4.A-B). In 2025, during recreational dives in Calpe (in the same area of the Penyal de Ifac), Dénia, Jávea and Moraira (Ampolla beach and surrounding waters) conducted by the authors of this study, the alga was also not detected. Taken together, these observations may indicate that its introduction in the area is likely to have occurred relatively recently.

The fact that a single specimen of *R. okamurae* has been discovered in shallow waters and subsequently remained absent during the three targeted follow-up dives inside a Marine Protected Area offers a very early starting point for management. This particular ZEC (Special Area of Conservation) of Calpe was established to conserve valuable Mediterranean habitats such as seagrass meadows, reefs, and the species that depend on them, ensuring their long-term ecological health. For the most part, at a visual level, this is being achieved. Thanks to the prohibition of destructive fishing methods, seabed disturbance, and anchoring in sensitive habitats, while low-impact recreation is generally allowed if it does not compromise conservation objectives. This has resulted in the area becoming a highly visited recreational diving area at a national level. Ultimately, with this early first detection of *R. okamurae*, the next steps should focus on involving local dive centers to raise awareness and leverage citizen science. Their cooperation will be essential for continuous non-invasive monitoring and early warning reporting to environmental authorities, rather than direct removal actions, to prevent accidental fragmentation and further clonal dispersal.

## CONCLUSIONS

The first record of *Rugulopteryx okamurae* in Calpe (Alicante) represents a remarkable event. Despite the limitations of this study, the detection of several specimens of *R. okamurae* in the area highlights its strong dispersal capacity, establishing the current northernmost distribution limit for the species in the Comunitat Valenciana. The absence of previous records in nearby areas during 2023–2025, combined with the finding of both drifting material in shallow waters and specimens recovered from commercial fishing nets, suggests a very recent front-wave colonization event extending across different depth ranges. This expansion is likely driven by marine currents or regional maritime activities, explicitly identifying the commercial fishing fleet as a major vector for secondary dispersal. Given that the species was recorded within a marine protected area and is already interacting with offshore fishing grounds, this presence highlights a potential ecological risk to vulnerable local benthic communities. Consequently, it is crucial that the competent authorities take these findings into consideration and implement a comprehensive spatiotemporal monitoring of the area and a dissemination and awareness campaign targeting diving clubs, fishermen, and other relevant stakeholders, with the aim of preventing the possible establishment of the species and supporting the adoption of preventive measures.

## ACKNOWLEDGEMENTS

The authors thank the Dirección General de Medio Natural y Animal (Conselleria de Medi Ambient, Infraestructures, Territori i Recuperació, GVA) and all the collaborators and field agents whose observations contributed to studying the distribution of this species in our waters.

Part of this research was carried out within the framework of the activities of the Marine Laboratory UA-Dénia, with the support of the Valencian Government (Generalitat Valenciana) and the Dénia City Council, and co-funded by the “Cátedra Interuniversitaria del Mar y la Sostenibilidad del Sector Náutico” promoted by Marina Port Valencia.

## AUTHORSHIP

E.R.G: Fieldwork, resources, writing original draft, writing – review & editing. J.F.: Fieldwork, resources, visualization, writing – review & editing. J.Y.D: Fieldwork, resources, writing – review & editing. E.S.F.: Fieldwork, resources, writing original draft, writing – review & editing. C.B: Funding acquisition, resources, writing – review & editing. C.P.M: Conceptualization, supervision, fieldwork, resources, taxonomic analysis, data curation, management, visualization, writing original draft, writing – review & editing.

